# Detection of F1 hybrids from single-genome data reveals frequent hybridization in Hymenoptera and particularly ants

**DOI:** 10.1101/2021.09.03.458849

**Authors:** Arthur Weyna, Lucille Bourouina, Nicolas Galtier, Jonathan Romiguier

**Author notes:** These authors share senior authorship (Nicolas Galtier, Jonathan Romiguier). First author. Last authors.

## Abstract

Hybridization occupies a central role in many fundamental evolutionary processes, such as speciation or adaptation. Yet, despite its pivotal importance in evolution, little is known about the actual prevalence and distribution of hybridization across the tree of life. Here we develop and implement a new statistical method enabling the detection of F1 hybrids from single-individual genome sequencing data. Using simulations and sequencing data from known hybrid systems, we first demonstrate the specificity of the method, and identify its statistical limits. Next, we showcase the method by applying it to available sequencing data from more than 1500 species of Arthropods, including Hymenoptera, Hemiptera, Coleoptera, Diptera and Archnida. Among these taxa, we find Hymenoptera, and especially ants, to display the highest number of candidate F1 hybrids, suggesting higher rates of recent hybridization in these groups. The prevalence of F1 hybrids was heterogeneously distributed across ants, with taxa including many candidates tending to harbor specific ecological and life history traits. This work shows how large-scale genomic comparative studies of recent hybridization can be implemented, uncovering the determinants of hybridization frequency across whole taxa.

## 1 Introduction

Hybridization, whereby members of genetically distinct populations mate and produce offspring of mixed ancestry (Abbott et al., 2013; Barton and Hewitt, 1985), has received much attention since the early days of evolutionary biology. From the onset, Darwin and his contemporaries spent a great deal of time studying hybrids and their fitness, which they recognized as a challenge to a discrete definition of species (Roberts, 1919). But the crucial importance of hybridization to biological evolution was fully realized only with the development of genetics in the following century. Formal studies of hybridization genetics led to the formulation of the biological species concept, and to the fundamental insight that speciation is generally driven by the evolution of isolating mechanisms in response to hybridization (Abbott et al., 2013; Dobzhansky, 1940; Mayr, 1942; Smadja and Butlin, 2011). The advent of genetic data also revealed the role of hybridization and introgression as important contributors to genetic variation and adaptation in many existing species (Anderson, 1953; Harrison and Larson, 2014), especially in the contexts of changing environments (Hamilton and Miller, 2016) and biological invasion (Prentis et al., 2008). Additionally, while hybridization was thought by many biologists to be relevant only for a few taxa such as plants (Barton, 2001), the accumulation of molecular data has continuously revealed its presence in many groups, including mammals, birds, fish, fungi, and insects (Taylor and Larson, 2019), with Mallet (2005) estimating that at least 10% of animal species frequently hybridize. These findings have further underlined the importance of hybridization in understanding many micro- and macro-evolutionary patterns across the tree of life (Abbott et al., 2013).

The same findings, however, also corroborated the old intuition that taxa can differ greatly in their susceptibility to hybridize, fueling discussions about the determinants of such heterogeneity (see Mallet, 2005 for a useful review). It was first understood that groups displaying a high number of sympatric species with low divergence, where the contact between compatible species is maximized, should be the most likely to hybridize (Edmands, 2002; Price and Bouvier, 2002). But sympatry and divergence are by themselves incomplete predictors of hybridization frequency, as strong reproductive barriers can arise from discrete evolutionary events (e.g., chromosome rearrangements or cytoplasmic incompatibilities; Bordenstein et al., 2001; Fishman et al., 2013), and can be rapidly selected for (i.e., reinforcement) or against depending on the relative fitness of hybrids (Smadja and Butlin, 2011). To understand heterogeneity in hybridization rates, is is thus important to also consider these ecological and phenotypic features of species that influence hybrid fitness, and more generally that influence the cost or gain in producing hybrids (Mallet, 2005). For instance, hybridization has been found to be more frequent in populations of spadefoot toads inhabiting ephemeral environments where hybrids outperform (Pfennig, 2007), or in rare species of birds where allospecific mates are easier to come by (Randler, 2002). A similar point was made by Mayr (1963), who suggested that polygamous species of birds should be the most likely to hybridize, because males with low parental investment should be more likely to accept interspecific mates. This early hypothesis is particularly significant in that it emphasizes on the idea that among characteristics of species relevant to hybridization, their life-history and mating system are of central importance.

One specific taxon in which relations between hybridization, mating systems and life-history have been extensively discussed is ants (Formicidae). Some ant genera are known to display unusually high rates of hybridization, based on both morphological and molecular data (Feldhaar et al., 2008; Nonacs, 2006; Umphrey, 2006). The first key trait of ants invoked to explain this pattern is haplodiploidy, a trait common to all Hymenoptera. Because males of Hymenoptera are haploids produced without fecundation, it is likely that hybrid sterility does not nullify the fitness of female Hymenoptera, which can still produce males after hybridizing (Feldhaar et al., 2008; Nonacs, 2006). This particularity of haplodiploids would hinder selection against hybridization and limit the formation of strict barriers to inter-specific mating. A second important ancestral trait of ants is eusociality, whereby reproductive females (i.e., queens) produce a large number of sterile helper individuals (i.e., workers) to form colonies. It was hypothesized that selection against hybridization is weaker in eusocial species because the fitness cost of hybrid sterility should be minimal in species producing a large majority of sterile individuals (Nonacs, 2006; Umphrey, 2006). This is especially likely in species in which queens mate multiply, and can combine inter- and intra-specific matings to ensure the production of a fraction of non-hybrid daughters (Cordonnier et al., 2020). Such interplay between hybridization, mating systems and life-history culminates in a handful of ant species that display unique hybridization-dependent reproductive systems, such as social hybridogenesis (Anderson et al., 2006; Fournier et al., 2005; Helms Cahan et al., 2002; Helms Cahan and Vinson, 2003; Kuhn et al., 2020; Lacy et al., 2019; Ohkawara et al., 2006; Pearcy et al., 2011; Romiguier et al., 2017). In these species, the cost of hybrid sterility is fully avoided because strong genetic caste determination constrains the development of hybrids towards the worker caste, while reproductive females can only be produced through intra-specific mating or parthenogenesis. The prevalence of hybridization-dependent systems within ants is virtually unknown (Anderson et al., 2008), but because they maintain large cohorts of F1-hybrid workers, they may help explain observations of high hybridization rates in ants.

While several hypotheses have been proposed to explain variation in hybridization rates across taxa, empirical comparative studies are still lacking, impeding any further understanding of its determinants. This is mainly due to the difficulties in evaluating the prevalence of hybridization at the group level. Methods to detect recent hybridization typically rely either on ambiguous morphological identification (which can lead to important ascertainment bias; Mallet, 2005), or on the use of large population-scale genetic samples including data for potential parental species (Anderson and Thompson, 2002; Payseur and Rieseberg, 2016; Schubert et al., 2017). These methods are sensitive and reliable in inferring hybrid status, but require large investments in time and money to produce results. To allow for comparative studies of hybridization at the level of entire taxa, it is necessary to implement methods that can be applied to many nonmodel species in parallel. In particular, methods applicable to the large volume of already published phylogenomic data (i.e., with one sequenced genome per species) would be especially desirable and cost-effective. For instance, phylogenomic data is available for more than 900 species of ants (223 represented genera), as the result of an extensive effort of Branstetter et al. (2017), who set a goal to sequence a large part of the diversity of Formicidae using standardized protocols (Faircloth et al., 2012). The same type of data has also been produced for many other Hymenoptera, and for other groups of Arthropods (including Hemiptera, Coleoptera, Diptera and Arachnida), paving the way for a comparative study of hybridization across these taxa of interest.

In this study, we implement a coalescent-based statistical method that allows for the detection of F1-hybrids using single diploid genomes. We first test this method and assess its efficiency using simulations and real data from identified F1 hybrid and non-hybrid individuals. We then apply the method to phylogenomic data, assessing the prevalence of F1 hybrids among five groups of Arthropods.

## 2 Materials and methods

### 2.1 Model

In this section we present the coalescent-based model of divergence which forms the basis of our F1 hybrid detection procedure. A F1 hybrid is the result of a cross between individuals from two different species. The heterozygosity of a such hybrid therefore reflects the divergence between its parental species, and can be modeled as shown in Figure 1. This model describes the expected distribution of the number of differences between two alleles found in a F1 hybrid in terms of two main parameters: the divergence time between the two parental species *t_s_* and the ancestral population size in their common ancestor *Ne* (fig. 1a). Briefly, if no migration occurred between the two parental lineages after their seperation, and if *t_s_* is large enough for lineage sorting to be complete, the total coalescence time of the two alleles is the sum of the divergence time *t_s_* and the coalescence time in the ancestral population *t_i_*. It is known from standard coalescence theory that the distribution of *t_i_* is well approximated by an exponential distribution with mean 2*Ne* (Wakeley, 2008). Consider a locus *i* of sequence length *l_i_* at which the two alleles of an F1 hybrid individual have been sequenced. Assuming an infinite-site mutation model with constant persite mutation rate *μ*, the number *n_i_* of observed allelic differences follows a Poisson distribution with mean 2*l_i_μ*(*t_i_* + *t_s_*), that is

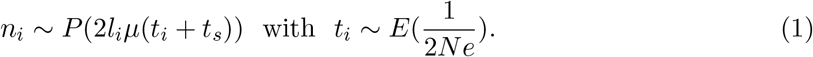

where *P* and *E* denote the Poisson and exponential distributions, respectively. Equation (1) leads to an expression for the probability to observe a number *k* of allelic differences between alleles at any given locus *i* in a F1 hybrid (see Appendix A for a complete derivation),

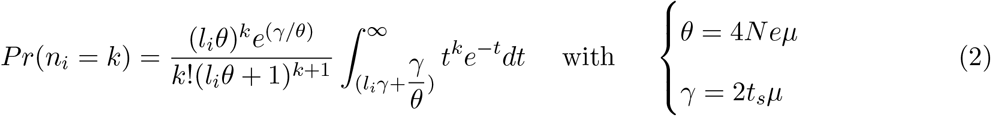

where *θ* is the ancestral population mutation rate, and *γ* is a measure of the heterozygosity acquired during the divergence process. Under the assumptions that *μ, t_s_* and *Ne* are constant across a set of *j* independent loci in a given diploid individual, each locus can be considered as a replicate of the same divergence scenario. In this case, the likelihood function of the set of observed numbers of differences between alleles is obtained by multiplying equation (2) across loci, and can be used to jointly estimate of *θ* and *γ*. To our knowledge this model was first introduced by Takahata et al. (1995), and later refined by Yang (1997), in the context of phylogenomics and ancestral population size estimation, with equation (2) being the continuous equivalent to equation (8) given in Yang (1997).

**Figure 1:**
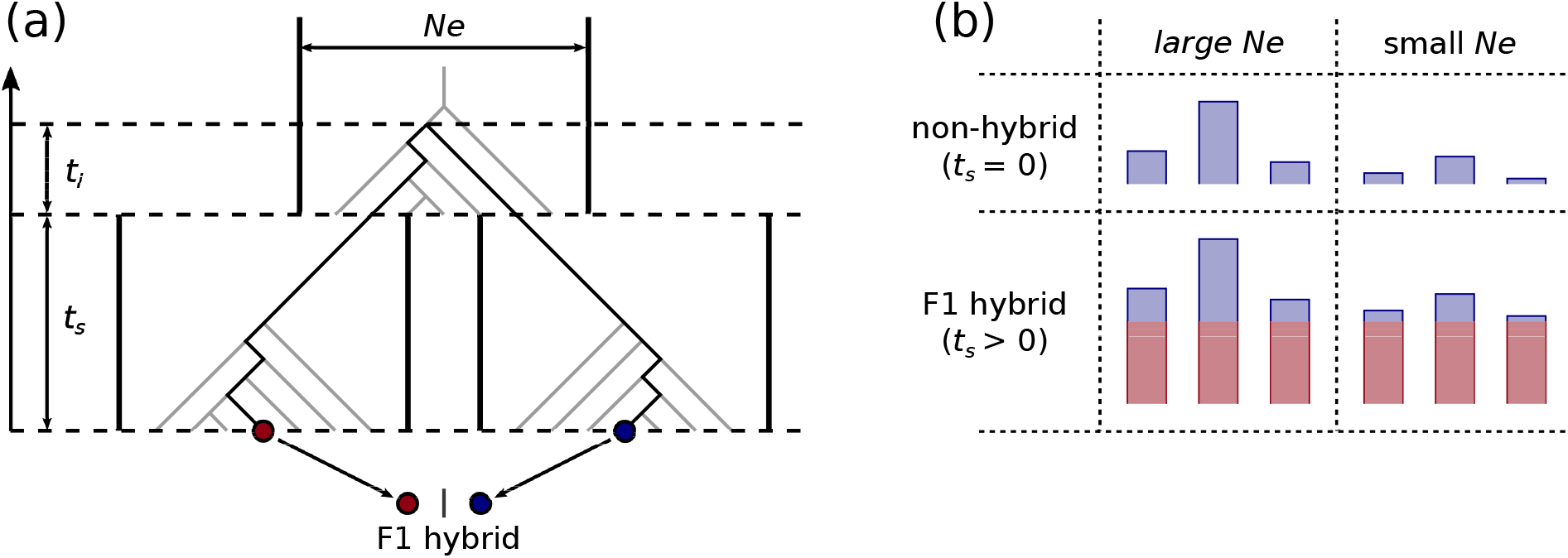
Coalescent-based model of divergence. (a) The population history assumed in the model of divergence described in the main text (eq. (1)). The darkened path represents the coalescence history of the two alleles (red and blue dots) that make up one locus in a diploid F1 hybrid. (b) Expected distribution of coalescence times for different values of *Ne* and *t_s_*. Blue bars represent the components of coalescence time linked to coalescence in the ancestral population. Red bars represent the uniform increase in coalescence times brought by divergence between parental populations.

Figure 1b illustrates the signal that is intended to be captured when estimating *γ* and *θ*. In a non-hybrid individual (i.e., whose parents belong to the same panmictic population), coalescence times between allele pairs are expected to follow an exponential distribution with mean and variance both determined by *Ne* (fig. 1b; top). In F1 hybrids, coalescence times are further increased by a fixed amount, which corresponds to the number of generations of divergence between the parental populations (1b; bottom, red bars). This uniform increase in coalescence times brought by divergence logically leads to an increased average coalescence time. This effect, however, is not by itself diagnostic of hybridization as it could be produced by an increase in *Ne*. Instead, what constitutes a unique signature of F1 hybrids is a decrease in the variance of coalescence times relative to the mean (fig. 1b, compare bottom to top). The relative variance in coalescence times is expected to be highest in non-hybrids, and to approach zero in F1 hybrids as the divergence between parental populations increases. The *γ* parameter captures this effect, whereas both *γ* and *θ* monitor the mean coalescence time. In other words, a non-zero estimate of *γ* means that the observed divergence between alleles is more similar across loci than expected under the standard coalescent.

Because the proposed statistical procedure partitions observed heterozygosity between *γ* and *θ*, it is expected that estimates of both parameters will be positively correlated with the genetic diversity of samples. For instance, a sample with low heterozygosity can only yield low estimates of *γ* and *θ*. For this reason, we mostly relied on the ratio *γ*/*θ*, which is not directly related to sample heterozygosity. This ratio is expected to be close to zero in non-hybrids, and non-zero in F1 hybrids. Furthermore, a *γ*/*θ* ratio above one implies that the divergence time between parental populations is longer than 2*Ne* generations, which is the expected time for complete lineage sorting. Such a high value is very unlikely to be reached by non-hybrid individuals.

### 2.2 Simulated test loci sets

To start evaluating our ability to detect F1 hybrids amongst diploid individuals, we simulated F1 hybrid, non-hybrid and 1^*st*^ generation backcross hybrid samples in the following manner.

Individual diploid loci were simulated by using *ms v2014.03.04* (Hudson, 2002) to sample pairs of alleles, together with the corresponding two-alleles gene trees, under the demographic scenario described in fig. 1a. To span across realistic values of both parameters of interest, values of *θ* and *γ* in simulations were set to be {10^−4^, 10^−3^, 10^−2^} and {0, 10^−4^, 10^−3^, 10^−2^}, respectively. Once obtained, gene trees were converted to explicit nucleotide sequences pairs through the application of a HKY mutation model using *seq-gen v1.3* (Rambaut and Grassly, 1997). The length of simulated sequences was set to be normally distributed with mean 1000bp and standard deviation 300bp. At this point, simulated F1 hybrid and non-hybrid individuals were constructed by putting together independent collections of sequences pairs simulated under *γ* > 0 and *γ* = 0, respectively. First generation backcross individuals were constructed by putting together both types of sequences pairs in random proportions following a binomial distribution with *p* = 0.5, (i.e., as expected from a backcross with random meiosis and no linkage). Ten individuals of each type were constructed for each possible combination of parameter values, and for two possible loci set sizes (200 or 500 loci). Finally, simulated individuals (i.e., loci sets) were sequenced in-silico using *art-illumina v2.5.8* (Huang et al., 2012). We emulated standard PE150 sequencing on HiSeq 2500 with 10X coverage, using a standard normally distributed fragment size with mean 400bp and standard deviation 20bp.

### 2.3 UCE datasets

In all applications to real data, we used ultra-conserved elements (UCEs) and their variable flanking regions as loci sets. UCEs are short (around 100bp on average) independent genomic regions that are conserved without duplication across large phylogenetic groups (Faircloth et al., 2012). These regions are usually sequenced through hybridization capture protocoles (Faircloth, 2017; Miles Zhang et al., 2019), but subsets of UCEs that correspond to transcribed genomic regions can also be retrieved from transcriptomic data (Bossert et al., 2019; Miles Zhang et al., 2019). This last fact is convenient in the context of this study, because transcriptome sequencing data is available for known hybrid systems, featuring *a priori* identified F1 hybrids and nonhybrid individuals, and can be used to further test our procedure using real data. We retrieved transcriptome sequencing data published on *genbank* from two types of well-characterized F1 hybrids: 12 hybrid workers from the harvester ant *Messor barbarus* (Romiguier et al., 2017), and 18 *Equus caballus x asinus* hybrids (9 mules and 9 hinnies; Wang et al., 2019). Data from the same sources for 7 haploid males and 5 non-hybrid queens of *M. barbarus*, as well as for 5 donkeys, were added for comparison. Genbank identifiers and metadata for *Messor* and *Equus* samples are available in supplementary tables S1 and S2, respectively.

Sequencing data obtained through UCE-capture protocoles has been published for a large number of non-model species, especially in Hymenoptera (Faircloth et al., 2012; Miles Zhang et al., 2019), thus allowing for a large scale search for F1 hybrids in these groups. We retrieved from *genbank* UCE-capture sequencing data from diploid samples belonging to groups of Arthropods for which specific capture probe sets were available: Formicidae (*“Insect Hymenoptera 2.5K version 2, Ant-Specific”* probe set; Branstetter et al., 2017), non-Formicidae Hymenoptera (“Insect Hymenoptera 2.5K version 2, Principal” probe set; Branstetter et al., 2017), Hemiptera (“Insect Hemiptera 2.7K version 1” probe set; Branstetter et al., 2017 and Kieran et al., 2018), Coleoptera (“Insect Coleoptera 1.1K version 1” probe set; Faircloth, 2017), Diptera (“Insect Diptera 2.7K version 1” probe set; Faircloth, 2017), and Arachnida (“Arachnida 1.1K version 1” probe set; Faircloth, 2017 and Starrett et al., 2016). To minimize the statistical weight of multiply sampled species, while maximizing statistical power at the group level, we kept only one sample per identified species (choosing samples with highest file size) and all samples lacking a complete identification (identified only to the genus level). Hymenoptera samples reported as males were considered as haploid and discarded. All remaining data files were downloaded from *genbank* using the *fasterq-dump* program from *SRA Toolkit v2.10.9*. Genbank identifiers and metadata for these samples are available in supplementary table S3.

### 2.4 Parameters estimation

To obtain estimates of *γ* and *θ* from simulated and real sequencing data, we systematically applied the following procedure. Raw read files were cleaned with *fastp v0.20.0* (Chen et al., 2018) to remove adapters, reads shorter than 40 bp, and reads with less than 70 percent of bases with a phred score below 20. Cleaned reads were then assembled using *megahit v1.1.3* (Li et al., 2015) with k-mer size spanning from 31 to 101 by steps of 10. The *phyluce v1.6* (Faircloth, 2016) tool suite (CIT) was used to identify and isolate UCE loci from de-novo assemblies, by blasting contigs against UCE probe sets with the *phyluce-assembly-match-contigs_to-probes* function. In this step, assemblies obtained from test samples of *M. barbarus* and *Equus* were blasted against the *“Insect Hymenoptera 2.5K version 2, Ant-Specific”* (Branstetter et al., 2017) and the *“Tetrapods 5K version 1”* (Faircloth et al., 2012) UCE probe sets, respectively. Likewise, assemblies obtained from UCE-capture samples were blasted against the probe set associated to their phylogenetic group. As no probe set exists for simulated loci, these were blasted against custom probe sets constructed from their true sequence (i.e., as output by *seqgen*). Following this step, cleaned sequencing reads were realigned to isolated loci using *bwa v0.7.17* (Li and Durbin, 2009) with default settings, and *angsd v0.921* (Korneliussen et al., 2014) was used to obtain allelic substitutions counts from read alignment files. Finally, we obtained estimates of *θ* and *γ* through bayesian estimations, using the *R* package *rstan v2.21.2* (Stan Development Team, 2019; Stan Development Team, 2020) and uninformative priors spanning all realistic values for both parameters (i.e., uniform priors constrained between 0 and 0.2). The mean of the posterior distribution of each parameter was used as a point estimate, while credibility intervals where constructed from its 2.5% and 97.5% quantiles.*R* scripts and the *stan* file necessary to run statistical estimations on a given set of observed allelic differences counts are available as supplementary documents (https://zenodo.org/record/5415947).

## 3 Results

### 3.1 Simulations

Applying our estimation procedure on simulated data, we find that our method can be used to efficiently discriminate F1 hybrids from non-hybrids and first generation backcross hybrids. Accurate divergence estimates can be obtained in simulated F1 hybrids using as little as 200 loci (fig. 2), provided that *γ* is in the same order of magnitude as the ancestral population mutation rate *θ* or higher (i.e., consistent with the model’s requirement of complete lineage sorting). Under the same condition, estimates of *γ* in non-hybrids and backcross hybrids are lower and do not exceed values one order of magnitude below estimated ancestral population size *θ* (which are themselves accurate; see fig. S1). Across all simulations, 61.1% of simulated F1 hybrids yielded both estimates of *γ* higher than 10^−3^ and estimates of *γ*/*θ* higher 1, while no backcross hybrids or non-hybrids did, demonstrating the specificity of the method. Furthermore, we find that increasing the number of sequenced loci from 200 to 500 does not increase our ability to identify F1 hybrids (see fig. S2), which suggest than 200 loci is a good minimal requirement in applications to real data.

**Figure 2:**
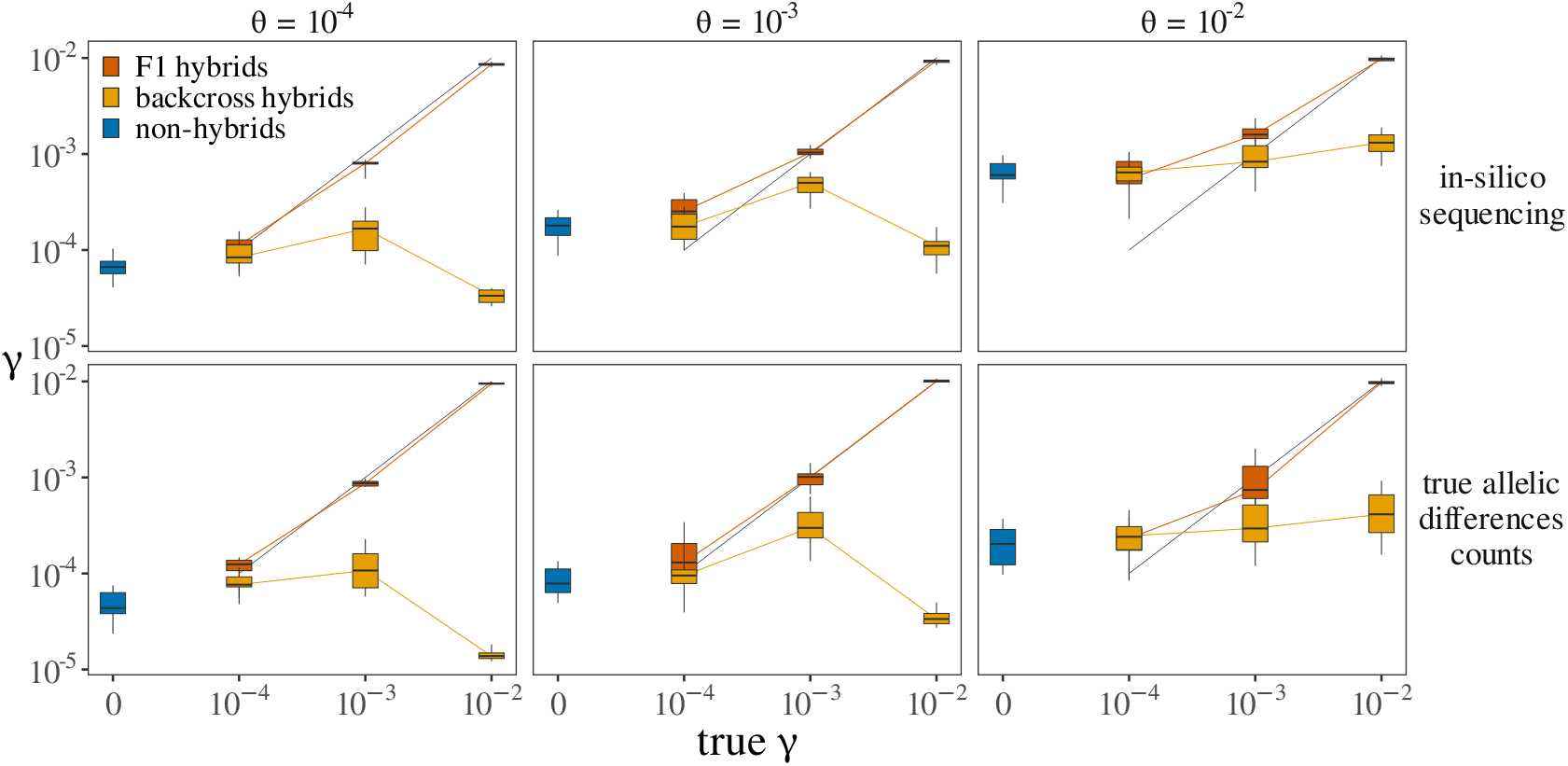
Estimates of divergence in simulated individuals. Each box represents the distribution of estimated *γ* values across 10 simulated individuals. Every individual consists of a collection of 200 loci simulated under a given combination of true *θ* (given in headers) and *γ* (given in x axis) values. In non-hybrid individuals *γ* is always zero. In backcross hybrids, the true value of *γ* is that given in the header, but only for a binomial proportion of loci (as described in the main text). The top row represents values obtained when estimating *γ* on loci sets obtained through the complete simulation procedure (including in-silico sequencing, read assembly and realignement, and substitutions counts estimation). The bottom row represent values obtained when estimating *γ* on sets of true counts data as output by ms (i.e., skipping subsequent simulation steps).

Simulations also revealed the statistical limits of our approach, which tends to overestimate the divergence parameter *γ* whenever the true ancestral population mutation rate *θ* is high (fig. 2, top row). This translates into estimates of *γ* departing from zero in non-hybrids with high overall polymorphism. Interestingly, this overestimation can be shown to arise in part from error in genome assembly, reads alignments, and estimations of allelic differences counts. When estimating parameters using sets of true allelic differences counts as first output by *ms* (fig. 2, bottom row), divergence overestimation is less important in hybrids and non-hybrids (fig. S2). This suggest that in real data, a positive correlation will be expected between divergence estimates and overall sample quality.

### 3.2 Accurate identification of F1 hybrids in two known hybrid systems

To further quantify our ability to distinguish between non-hybrids and typical F1 hybrid individuals, we applied our estimation procedure to sequencing data from two types of well-characterized F1 hybrids, hybrid workers from the harvester ant *Messor barbarus* (Romiguier et al., 2017), and *Equus caballus x asinus* hybrids (mules and hinnies) (Wang et al., 2019). Sequencing data from the same sources for males and non-hybrid queens of *M. barbarus*, as well as for donkeys, were added to the analysis for comparison. This analysis confirmed that F1 hybrids and non-hybrid individuals can be discriminated without ambiguity (fig. 3; parameters estimates are given in table S1 and S2). Estimates of divergence (*γ*) in F1 hybrids always strongly departed from 0 and showed little variation across samples (3.39 × 10^−3^ ± 2.05 × 10^−4^ sd in *M. barbarus* workers; 1.47 × 10^−3^ ± 1.46 × 10^−4^ sd in mules and hinnies). By contrast, estimated values of *γ* in nonhybrid samples were always much closer to 0 in non-hybrid individuals (2.34 × 10^−5^ ±3.29 × 10^−5^ sd in *M. barbarus* males and queens; 1.63 × 10^−5^ ± 3.31 × 10^−6^ sd in donkeys). The ratio *γ*/*θ* reached the critical value of one in *M. barbarus* workers (1.067 ± 0.190 sd) while being two orders of magnitude lower in males and queens (0.012 ± 0.010 sd). This confirms that such a threshold value is reliable for discriminating true F1 hybrids. Ratios obtained in mules and hinnies are lower than 1 however (0.418 ± 0.061 sd), further suggesting that *γ*/*θ* > 1 is a conservative requirement likely not to be reached by many true F1 hybrid. Interestingly, UCE-capture data for a single worker of *M. barbarus* (fig. 3a) led to slightly higher parameter estimates than transcriptomic data, but to a similar *γ*/*θ* ratio (1.131). This suggests that UCEs that could be retrieved from transcriptomes of *M. barbarus* are less polymorphic on average, but contain the same information regarding relative divergence and hybrid status.

**Figure 3:**
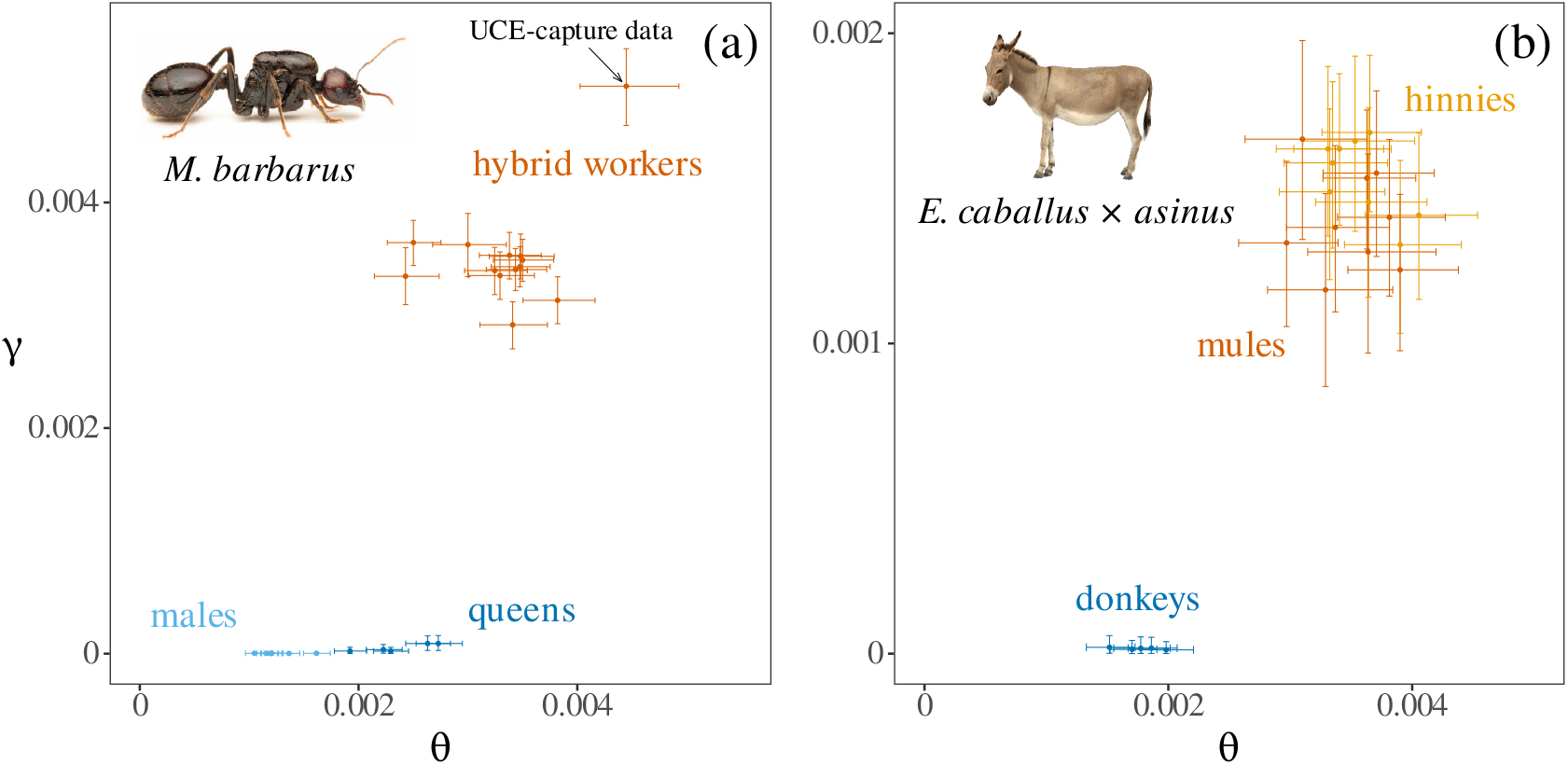
Discrimination of F1 hybrids in transcriptomes of *M. barbarus* and *Equus*. Estimated values for the divergence parameter *γ* and the ancestral population mutation rate *θ* are represented for *M. barbarus* (a) and *Equus* (b). Colored points and lines represent point estimates and confidence intervals, respectively. Values obtained using UCE-capture data for a single worker of *M. barbarus* (genbank:SRR5437981) were added for comparison (arrow in panel a).

### 3.3 High prevalence of F1 hybrids in Hymenoptera and Formicidae

The application of our procedure to UCE-capture data, comprised of many samples of heterogeneous quality, led to the observation of the quality bias predicted from simulations. Specifically, we noted that older samples yielded slightly higher *γ*/*θ* estimates on average than recent ones, resulting in a significant correlation between the later ratio and specimen collection date (*ρ* = −0.275, p-value < 2.2 × 10^16^). This bias is most likely due to lower sequence quality and increased data treatment error in old specimen, which leads to an overestimation of *γ* as mentioned in the previous section. To take this effect into account, we excluded samples collected before 1980 and specimen with unknown collection date from subsequent analyses. This does not eliminate the mentioned correlation which remains significant (*ρ* = −0.163, p-value = 3.43 × 10^−11^), but ensures that no old, highly degraded sample is wrongly interpreted as a hybrid. This choice of a threshold date does not affect our subsequent statistical results (see table S4). We also discarded samples for which less than 200 UCE locis could be retrieved to ensure sufficient statistical power. After application of these filters, we could obtain parameter estimates (figure 4) for 850 Formicidae (223 represented genera), 472 other Hymenoptera (288 genera), 177 Hemiptera (121 genera), 51 Coleoptera (45 genera), 25 Diptera (5 genera) and 65 Arachnida (56 genera). All parameter estimates can be found in supplementary table S3.

Our results revealed important differences between phylogenetic groups regarding the prevalence of F1 hybrids. We found several candidate F1 hybrids (*γ*/*θ* > 1) in Formicidae (29 candidates; figure 4a) and other Hymenoptera (15 candidates; figure 4b), while none were found in Hemiptera, Coleoptera, Diptera (figure 4c) or Arachnida (figure 4d). This result can not be explained by the larger number of Hymenoptera available, as under the observed frequency of candidates in this group (0.033), the probability to observe no candidates in other groups would be 8.36 × 10^−5^. Species names, divergence estimates and metadata for all candidate F1 hybrids can be found in table 1. In Formicidae, two samples originating from species known to produce F1 hybrid workers (*M. barbarus* and *Wasmannia Auropunctata*) were identified as candidate F1 hybrids, further validating our method. Interestingly, a third known F1 hybrid (*Paratrechina longicornis*) was found to fall below the required value of *γ*/*θ* > 1, again showing that this value is conservative and likely to produce many false negatives. Beyond individual candidates, Formicidae also displayed a significantly higher mean *γ*/*θ* ratio than non-Hymenoptera insects, as evidence by a post-hoc Tukey honest significance test (figure 4e). This suggests that, on average, successful interspecific mating is more frequent in ants than in other groups. Finally, candidates F1 hybrids displayed higher average divergence *γ* in Formicidae than in other Hymenoptera (T = 2.24, p-value = 0.0324), suggesting that hybridization events in this group tend to occur between more divergent individuals. Note that these two last results are mostly unchanged under other reasonable choices of threshold dates (see table S4).

**Figure 4:**
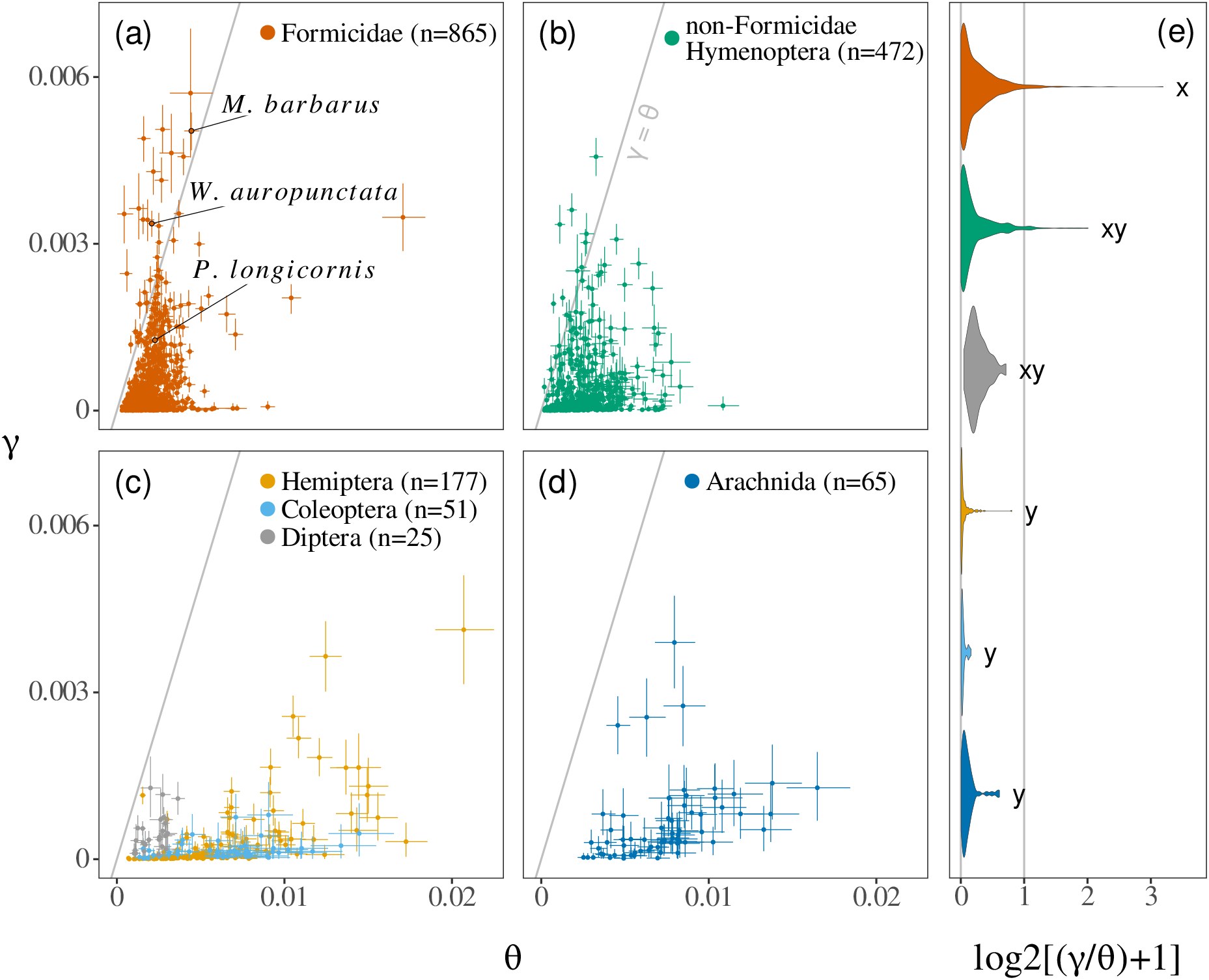
Genomic scans for hybridization in six groups of arthropodes. Estimates of the divergence parameter *γ* and the ancestral population mutation rate *θ* are represented for Formicidae (a), Non-Formicidae Hymenoptera (b), other insects (c) and Arachnida (d). Colored points and lines represent bayesian point estimates and credibility intervals (see main text), respectively. (e) Distribution of the ratio *γ*/*θ* in each group. A one-shifted log2-scale, under which the critical value of *γ*/*θ* = 1 is unchanged, was used for visual convenience. Letters summarize the result of a post-hoc Tukey honest significance test, carried out using the *HSD.test* function of the *R* package *agricolae* (CIT). Groups with no letters in common have significantly different means (with *α* = 0.05). All results were obtained using only dated and recent samples (see main text).

**Table 1:**
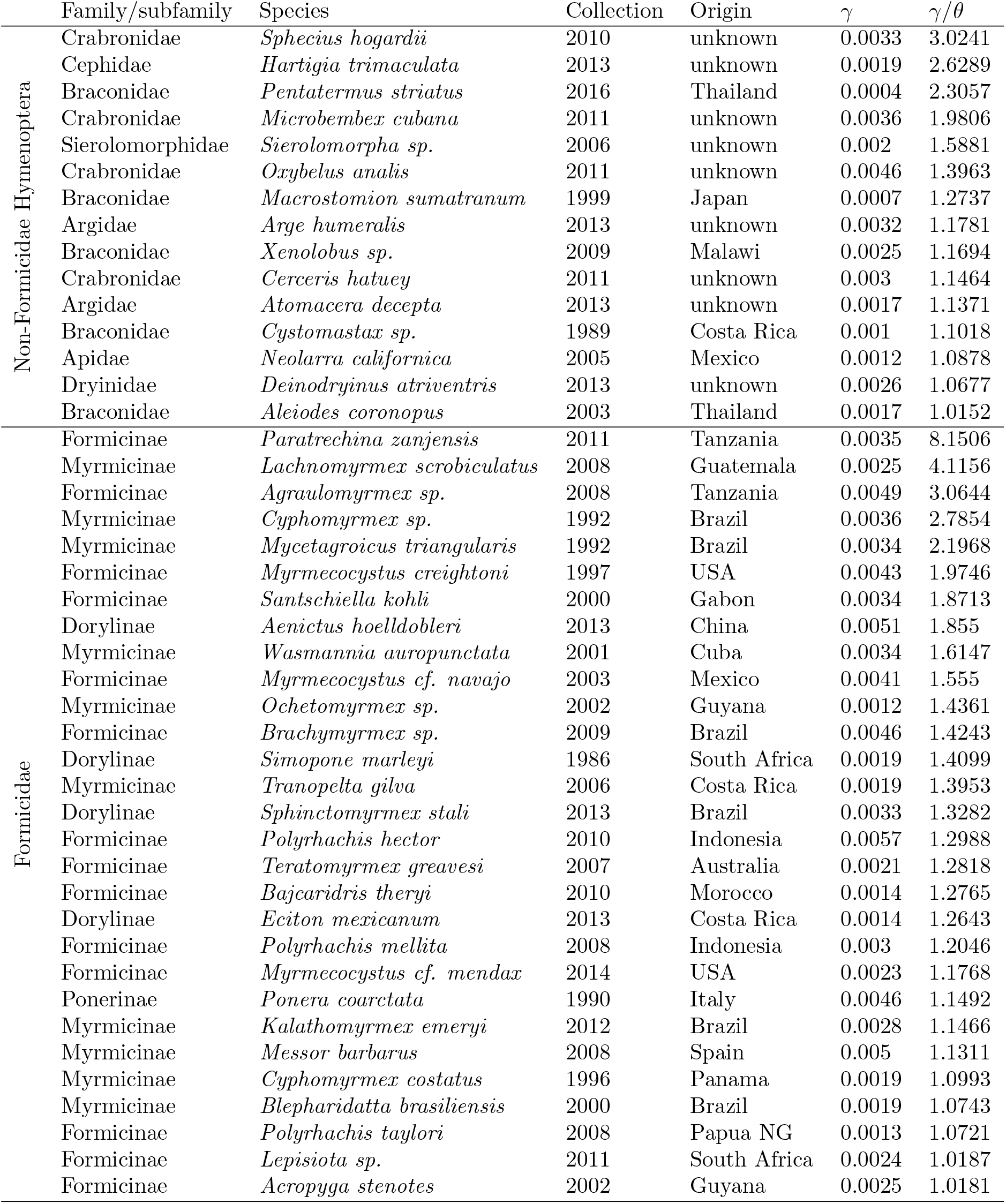
Candidate F1 hybrids. The table gives metadata and point parameter estimates for each candidate F1 hybrid (i.e. *γ*/*θ* > 1) in our analysis.

| | Family/subfamily | Species | Collection | Origin | $\gamma$ | $\gamma/\theta$ |
| --- | --- | --- | --- | --- | --- | --- |
| Non-Formicidae Hymenoptera | Crabronidae | *Sphecius hogardii* | 2010 | unknown | 0.0033 | 3.0241 |
|  | Cephididae | *Hartigia trimaculata* | 2013 | unknown | 0.0019 | 2.6289 |
|  | Braconidae | *Pentatermus striatus* | 2016 | Thailand | 0.0004 | 2.3057 |
|  | Crabronidae | *Microbembex cubana* | 2011 | unknown | 0.0036 | 1.9806 |
|  | Sierolomorphidae | *Sierolomorpha sp.* | 2006 | unknown | 0.002 | 1.5881 |
|  | Crabronidae | *Oxybelus analis* | 2011 | unknown | 0.0046 | 1.3963 |
|  | Braconidae | *Macrostomion sumatranum* | 1999 | Japan | 0.0007 | 1.2737 |
|  | Argidae | *Arge humeralis* | 2013 | unknown | 0.0032 | 1.1781 |
|  | Braconidae | *Xenolobus sp.* | 2009 | Malawi | 0.0025 | 1.1694 |
|  | Crabronidae | *Cerceris hatuey* | 2011 | unknown | 0.003 | 1.1464 |
|  | Argidae | *Atomacera decepta* | 2013 | unknown | 0.0017 | 1.1371 |
|  | Braconidae | *Cystomastax sp.* | 1989 | Costa Rica | 0.001 | 1.1018 |
|  | Apidae | *Neolarra californica* | 2005 | Mexico | 0.0012 | 1.0878 |
|  | Dryinidae | *Deinodryinus atriventris* | 2013 | unknown | 0.0026 | 1.0677 |
|  | Braconidae | *Aleiodes coronopus* | 2003 | Thailand | 0.0017 | 1.0152 |
| Formicidae | Formicinae | *Paratrechina zanzensis* | 2011 | Tanzania | 0.0035 | 8.1506 |
|  | Myrmicinae | *Lachnomyrmex scrobiculatus* | 2008 | Guatemala | 0.0025 | 4.1156 |
|  | Formicinae | *Agraulomyrmex sp.* | 2008 | Tanzania | 0.0049 | 3.0644 |
|  | Myrmicinae | *Cyphomyrmex sp.* | 1992 | Brazil | 0.0036 | 2.7854 |
|  | Myrmicinae | *Mycetagroicus triangularis* | 1992 | Brazil | 0.0034 | 2.1968 |
|  | Formicinae | *Myrmecocystus creightoni* | 1997 | USA | 0.0043 | 1.9746 |
|  | Formicinae | *Santschiella kohli* | 2000 | Gabon | 0.0034 | 1.8713 |
|  | Dorylinae | *Aenictus hoelldobleri* | 2013 | China | 0.0051 | 1.855 |
|  | Myrmicinae | *Wasmannia auropunctata* | 2001 | Cuba | 0.0034 | 1.6147 |
|  | Formicinae | *Myrmecocystus cf. navajo* | 2003 | Mexico | 0.0041 | 1.555 |
|  | Myrmicinae | *Ochetomyrmex sp.* | 2002 | Guyana | 0.0012 | 1.4361 |
|  | Formicinae | *Brachymyrmex sp.* | 2009 | Brazil | 0.0046 | 1.4243 |
|  | Dorylinae | *Simopone marleyi* | 1986 | South Africa | 0.0019 | 1.4099 |
|  | Myrmicinae | *Tranopelta gilva* | 2006 | Costa Rica | 0.0019 | 1.3953 |
|  | Dorylinae | *Sphinctomyrmex stali* | 2013 | Brazil | 0.0033 | 1.3282 |
|  | Formicinae | *Polyrhachis hector* | 2010 | Indonesia | 0.0057 | 1.2988 |
|  | Formicinae | *Teratomyrmex greavesi* | 2007 | Australia | 0.0021 | 1.2818 |
|  | Formicinae | *Bajcaridris theryi* | 2010 | Morocco | 0.0014 | 1.2765 |
|  | Dorylinae | *Eciton mexicanum* | 2013 | Costa Rica | 0.0014 | 1.2643 |
|  | Formicinae | *Polyrhachis mellita* | 2008 | Indonesia | 0.003 | 1.2046 |
|  | Formicinae | *Myrmecocystus cf. mendax* | 2014 | USA | 0.0023 | 1.1768 |
|  | Ponerinae | *Ponera coarctata* | 1990 | Italy | 0.0046 | 1.1492 |
|  | Myrmicinae | *Kalathomyrmex emeryi* | 2012 | Brazil | 0.0028 | 1.1466 |
|  | Myrmicinae | *Messor barbarus* | 2008 | Spain | 0.005 | 1.1311 |
|  | Myrmicinae | *Cyphomyrmex costatus* | 1996 | Panama | 0.0019 | 1.0993 |
|  | Myrmicinae | *Blepharidatta brasiliensis* | 2000 | Brazil | 0.0019 | 1.0743 |
|  | Formicinae | *Polyrhachis taylori* | 2008 | Papua NG | 0.0013 | 1.0721 |
|  | Formicinae | *Lepisiota sp.* | 2011 | South Africa | 0.0024 | 1.0187 |
|  | Formicinae | *Acropyga stenotes* | 2002 | Guyana | 0.0025 | 1.0181 |

Samples for roughly two thirds (223 represented genera) of the diversity of ants (about 300 genera, Bolton) genera were available for this study. This allowed us to evaluate whether the distribution of hybridization within ants genera is random. Positioning candidate F1 hybrids on a phylogeny of ants genera (figure 5) and the application of Abouheif’s test (Abouheif, 1999) revealed a significant positive phylogenetic correlation in mean *γ*/*θ* across genera (C = 0.1758; p-value = 0.007). This can be explained by the absence of candidate F1 hybrids from widely sampled groups, such as the Crematogastrini tribe (171 species from 59 genera), and by their high prevalence in other groups, such as the Attini tribe (10 candidates representing 9.4% of the tribe’s sampled species). Genera *Cyphomyrmex, Polyrhachis* and *Myrmecocystus* also displayed several distinct candidate F1 hybrids.

**Figure 5:**
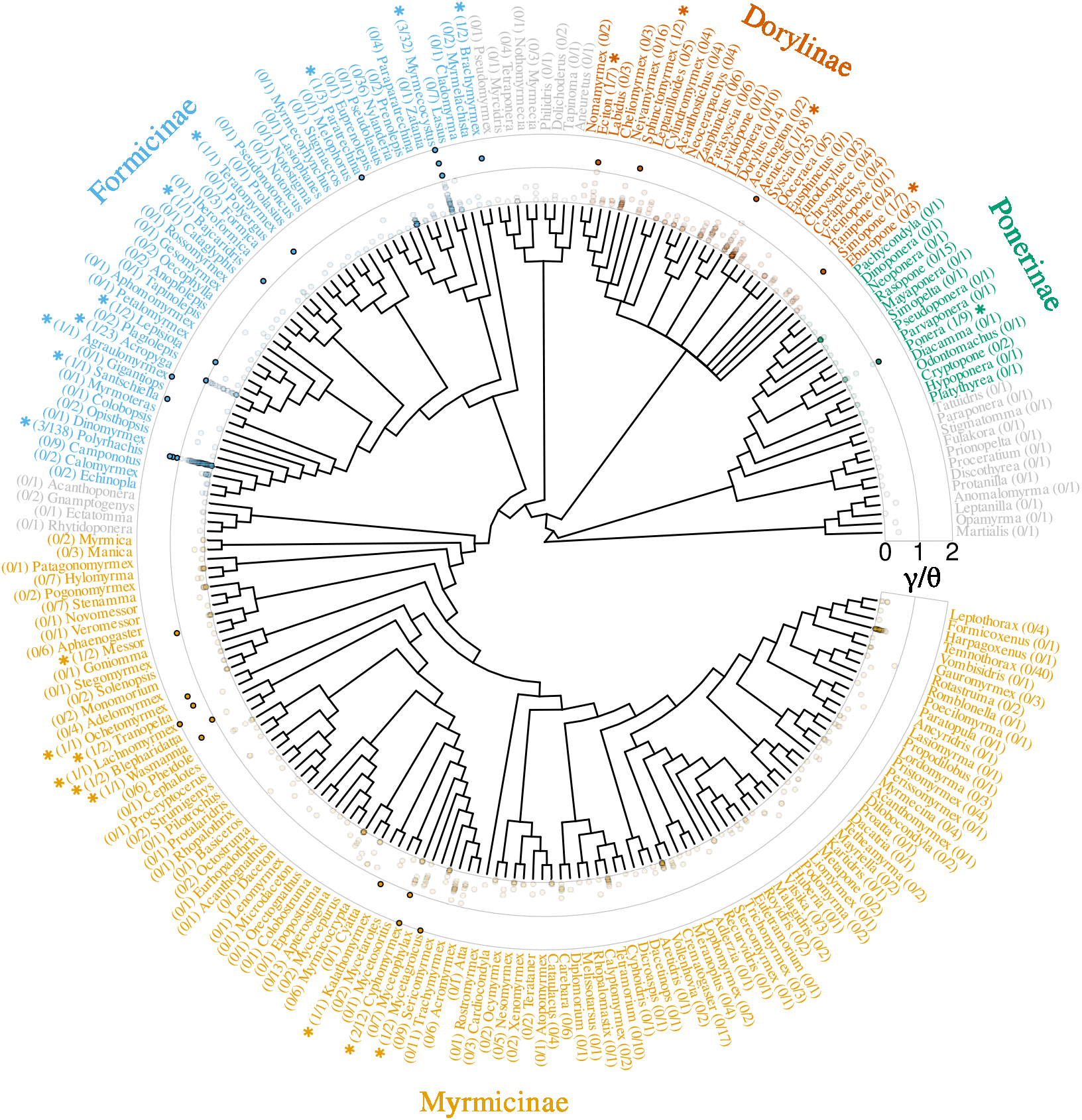
Occurence of F1 hybrids across genera of Formicidae. Estimates of the ratio *γ*/*θ* obtained in Formicidae are represented against the topology of genera in this group (retrieved from Antwiki). Genera counting at least one species with *γ*/*θ* > 1 (i.e., probable F1 hybrid) are highlighted by a star. The number of such species per genera, as well as the total number of species per genera are given for each genus. *γ*/*θ* ratios higher than 2 were truncated to 2 for readability. Three genera with no candidate F1 hybrids (*Cryptopone, Pseudoatta* and *Strongylognathus*), present in UCE capture data but not in the present topology, were not integrated in this representation or in statistical test for phylogenetic correlation.

## 4 Discussion

### 4.1 F1 hybrid detection from single genomes

In this article we implement and showcase a fast and flexible statistical method for F1 hybrids detection. This method only relies on the distribution of heterozygosity across a set of diploid loci, and is thus theoretically applicable to any type of polymorphic loci set such as UCE loci, coding genes or even RAD tags. Note however that chosen loci should ideally be 1-1 orthologs, in order to limit the risk of paralogy inflating observed substitutions counts and facilitate intragroup estimates comparisons. Besides its applicability to a large range of data types, the method is also flexible in that it does not rely on the use of parental genomes, unlike population-centered hybrid detection approaches (Anderson and Thompson, 2002; Payseur and Rieseberg, 2016; Schubert et al., 2017). It is thus especially suited for preliminary hybrid status assessment in single-species datasets composed of many non-model species (i.e., most phylogenomic datasets). In addition to applications in the study of hybridization prevalence across taxa (i.e., as done in this study), F1 hybrid detection could help preventing the use of error-inducing hybrids nuclear genomes in reconstructing species trees (McDade, 1992).

Naturally, the presented method also has some shortcomings, which stem in the limited statistical power provided by single individual genomes. Perhaps the most important limitation of the method is that it is restricted to the discrimination of F1 hybrids, and can not reliably be used to identify backcross hybrids. This restricts the use of the method to the study of present hybridization and suggests that many hybrids can be missed, given the rarity of true F1 hybrids in natural populations. We have also shown that statistical error inherent to data treatment can inflate divergence estimates and lead to false identification of F1 hybrids. This is because our method relies on the assumption that divergence is characterized by a uniform increase in heterozygosity across loci. As sequencing, assembly and gene identification errors are likely to produce such an increase, their effect is mostly indistinguishable from true divergence using single genomes. This limits the application of our method to samples of good quality and limits its ability to identify F1 hybrids with low overall polymorphism. The sensitivity of the method is also hindered by any violation of the hypothesis of constant mutation rate in time and across loci. In fact, Yang (1997) has shown that variation in mutation rates generally reduces estimates of divergence by eroding any uniform component of heterozygosity. The limited sensitivity of the method might be problematic in other settings, but acts as a safeguard in our case by making F1 hybrids detection more conservative.

### 4.2 High prevalence of F1 hybrids in Hymenoptera and particularily in ants

F1 hybrids detection in 850 Formicidae, 472 non-Formicidae Hymenoptera, 177 Hemiptera, 51 Coleoptera, 25 Diptera and 65 Arachnida revealed a heterogeneous distribution of F1 hybrids prevalence across these groups. We identified 29 and 15 candidate F1 hybrids in Formicicidae and other Hymenoptera, respectively, and none in other groups, a result that can not be explained by uneven group sampling. High hybridization rates in Hymenoptera have been predicted by other authors (Feldhaar et al., 2008; Nonacs, 2006) under the rationale that haplodiploidy could mitigate the potential costs of out-breeding. More specifically, it was proposed that because haplodiploid females produce part of their descendants asexually, they should retain positive fitness even when engaging in non-viable inter-specific mating, leading to a weaker long-term selection against such behavior. While our results are compatible with this hypothesis, similar analyses on haplodiploid groups other than Hymenoptera will be necessary to confirm that haplodiploidy is the only factor explaining this pattern.

Within Hymenoptera, our analyses also revealed a significantly higher prevalence of F1 hybrids in Formicidae than in other Hymenoptera. High hybridization rates were previously described in a several ant genera (e.g. in some North American *Solenopsis* or European *Temnothorax*, Feldhaar et al., 2008), and have been suspected to be frequent in ants in general on the basis of several arguments. Some authors have hypothesized that hybrid sterility has a minimal fitness cost in eusocial species because they produce a large majority of normally sterile individuals (i.e., workers), leading to weaker selection against hybridization (Nonacs, 2006; Umphrey, 2006). The same authors also proposed that eusocial queens could use inter-specific mating as a “best of a bad situation” strategy allowing for the production of a workforce and the successful rearing of haploid sons in the absence of conspecific mates (e.g. in locally rare species). Such strategy, sometimes referred to as “sperm parasitism”, would be especially likely to arise if hybrid workers outperform regular ones, a hypothesis for which empirical evidence is still lacking (Feldhaar et al., 2008; Julian and Cahan, 2006; Ross and Robertson, 1990 but see James et al., 2002). Interestingly, the idea that eusociality facilitates or promotes hybridization is not clearly supported by the present analysis, as no candidate F1 hybrids were identified amongst 66 available non-Formicidae eusocial species (22 represented genera). While this might be because most of these species display relatively simple forms of eusociality as compared to ants (with 44 species belonging to either *Lasioglossum* or *Bombus*), it could also indicate that ants possess other traits relevant to frequent hybridization. Among characteristics unique to ants, the extreme functional simplification of workers (Peeters and Ito, 2015) could have favored hybridization by making hybrid individuals less affected by inherent developmental defects (e.g., fluctuating asymmetry). Additionally, the typically low morphological and behavioral divergence observed between males of related ant species has been proposed to reduce pre-mating barriers to hybridization in this group (Feldhaar et al., 2008).

### 4.3 Phylogenetic and ecological characteristics of F1 hybrids in ants

Beyond the higher prevalence of candidate F1 hybrids in ants, our analysis reveals that their phylogenetic distribution in the group follows a non-random pattern, hinting towards a potential connection with variation in ecological and life-history characteristics of species. One peculiar characteristic of some ants that is especially relevant to our findings is their display of hybridization-dependent reproductive systems. In these systems, strong genetic caste determination enforces that all workers are F1 hybrids developing from eggs fertilized by allospecific males (i.e., social hybridogenesis, as in *Messor, Pogonomyrmex, Solenopsis* or *Cataglyphis*; Anderson et al., 2006; Helms Cahan et al., 2002; Helms Cahan and Vinson, 2003; Kuhn et al., 2020; Lacy et al., 2019; Romiguier et al., 2017) or by males from a divergent lineage of the same species (i.e., as in *Wasmannia auropunctata, Vollenhovia emeyri* or *Paratrechina longicornis*; Fournier et al., 2005; Ohkawara et al., 2006; Pearcy et al., 2011), while queens are produced through regular intra-lineage mating or thelytokous parthenogenesis. In genera where it has been described, strong genetic caste determination has typically evolved independently multiple times (Anderson et al., 2006; Kuhn et al., 2020; Romiguier et al., 2017), indicating that phylogenetic correlation in this trait is expected. Furthermore, out of the three available species known to display such system (*M. barbarus* and *W. auropunctata*), two clearly stand out as F1 hybrids, indicating that our method is able to detect the divergence signal present in individual genomes of their workers. Overall, this suggests that other candidate F1 hybrids identified in this work might belong to species with similar reproductive systems, which would help explain why we detected a larger proportion of F1-hybrids in ants. This possibility echoes the prediction of some authors that the prevalence of strong genetic caste determination in Formicidae might have been largely underestimated (Anderson et al., 2008). Note for instance the case of the *Paratrechina zanjensis*, which displays the highest divergence signal within the analyzed data (tab. 1) and belongs to the same genus as *P. longicornis*, a species known to produce hybrid workers (Pearcy et al., 2011). Further investigation of the population genetics of *P. zanjensis* might reveal a system similar to that found in *P. longicornis*, but with a longer history of divergence.

If detected candidates correspond to undetected cases of strong genetic caste determination, our results might help shed new light on the conditions that drive the evolution of such systems. For instance, it has been hypothesized that genetic caste determination evolves more frequently in taxa with a highly specialized diet (such as granivory), as a reduced dietary spectrum would impede the use of differential larval feeding as a mean to drive caste determination (Romiguier et al., 2017). Interestingly, we found significantly higher *γ*/*θ* ratios in genera listed as strictly herbivorous (fungus-growing, granivorous or specialized aphid-rearing diets; Blanchard and Moreau, 2016) than in omnivorous or carnivorous genera (one-sided Welsch t-test; t = 3.3292, df = 154.75, p-value = 0.00054). This may suggest that highly specialized diets do favor the evolution of genetic caste determination. This remains highly speculative however without an extended study on more genera and clear confirmation that *γ*/*θ* variations are mainly due to unusual reproductive systems across ants. While the exact proportion of detected F1-hybrids that are due to such reproductive systems is unknown at this stage, species with *γ*/*θ* ratios superior to known cases (*M. barbarus, W. auropunctata, P. longicornis*, see Fig 4) would be good first candidates for future studies.

Besides unusual reproductive systems, high hybridization rates in Dorylinae and in Attini could be linked to the unusually high polyandry observed in these group (Keller and Reeve, 1994; Strassmann, 2001). Queens that mate multiply are less likely to mate only with interspecific males (Umphrey, 2006), and are therefore expected to display lower pre-mating barriers to hybridization. Such effect of polyandry is especially likely when both types of males are easily accessible, as in species with massive mating flights that are synchronized with other sympatric species. Such pattern is more frequent in species inhabiting temperate and arid climates, where mating flights are often triggered by heavy rainfall (Dunn et al., 2007). In favor of such connection, we find that the previously unsuspected xerophile genus *Myrmecocystus* counts several candidate F1 hybrids. As a final remark, we note that some ant groups display a high proportion of candidate F1 hybrids, while presenting no obvious life-history or ecological features likely to produce such pattern. This is especially true of the paraphyletic group of attines composed of *Ochetomyrmex, Tranopelta, Lachnomyrmex, Blepharidatta* and *Wasmannia*. This suggests the existence of other unknown factors in species predisposition to hybridization, and new biological models for the study of such factors.

## 5 Conclusion

Hybridization is a widespread and fundamental phenomenon that carries implications for many central processes of biological evolution, including speciation and adaptation. Here we present the first large-scale comparative study of hybridization prevalence in Arthropods, analyzing genomic data for more than 1500 non-model species obtained from public repositories. We report high rates of recent hybridization in Hymenoptera, and especially in ants, confirming previous predictions found in the literature. We also find the prevalence of F1 hybrids to be heterogeneously distributed within ants, with probable links with ecological and life-history features. These results were produced through the implementation of a scalable F1 hybrids detection method, which is applicable to virtually any modern sequencing data. Further applications of this method should help better assessing the frequency of hybridization across the tree of life, and understanding its determinants.

## Statement of authorship

AW and JR conceived the study. AW and NG developed statistical methods. LB and JR preformed preliminary analyses. AW performed the final analysis and wrote the first draft of the manuscript under the guidance of JR and NG. All authors contributed to the final version.

## Data accessibility

Supplementary tables containing all results produced in this work, as well as scripts and files necessary to apply our statistical procedure, are available here: https://zenodo.org/record/5415947.

## Appendices

## A Complete model derivation

With *T* = *t_s_* + *t_i_* the total coalescence time between two divergent alleles (see fig. 1), the probability to observe a number *k* of differences between them is

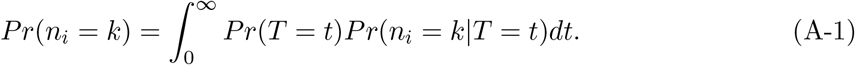

As the total coalescence time between two divergent alleles has to be greater than or equal to the divergence time *t_s_*, and as *t_i_* is exponentially distributed with mean 2*Ne*, we can write

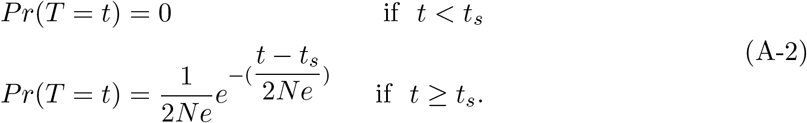

Assuming a infinite-site mutation model with constant per-site mutation rate *μ*, the number *n_i_* of expected substitutions between two alleles that diverged for *t* generations follows a Poisson distribution with mean 2*l_i_μt*, that is

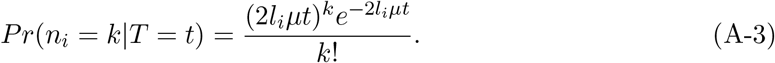

Plugging equations A-2 and A-3 into equation A-1 leads to equation 2 presented in the main tex

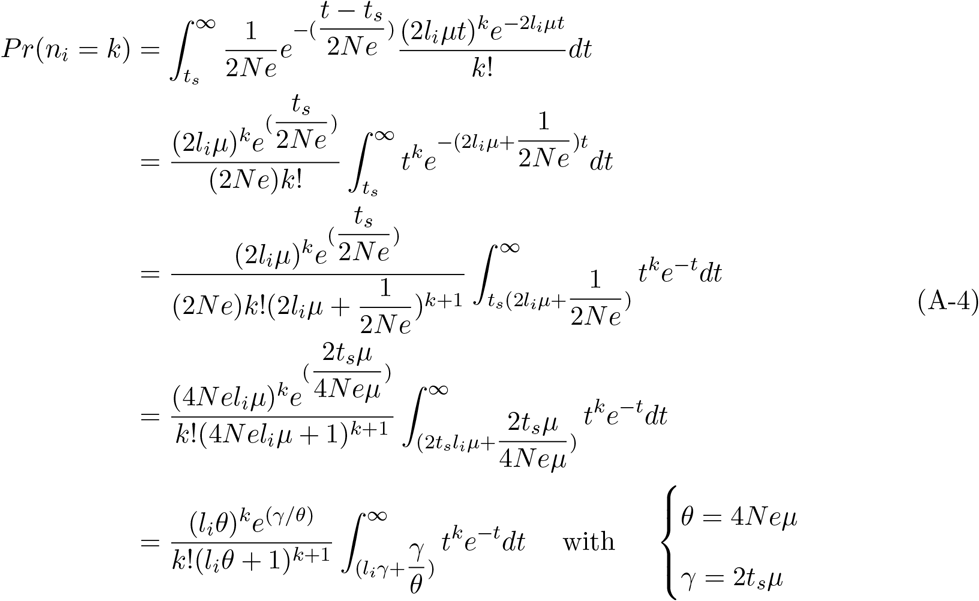

**Figure S1:**
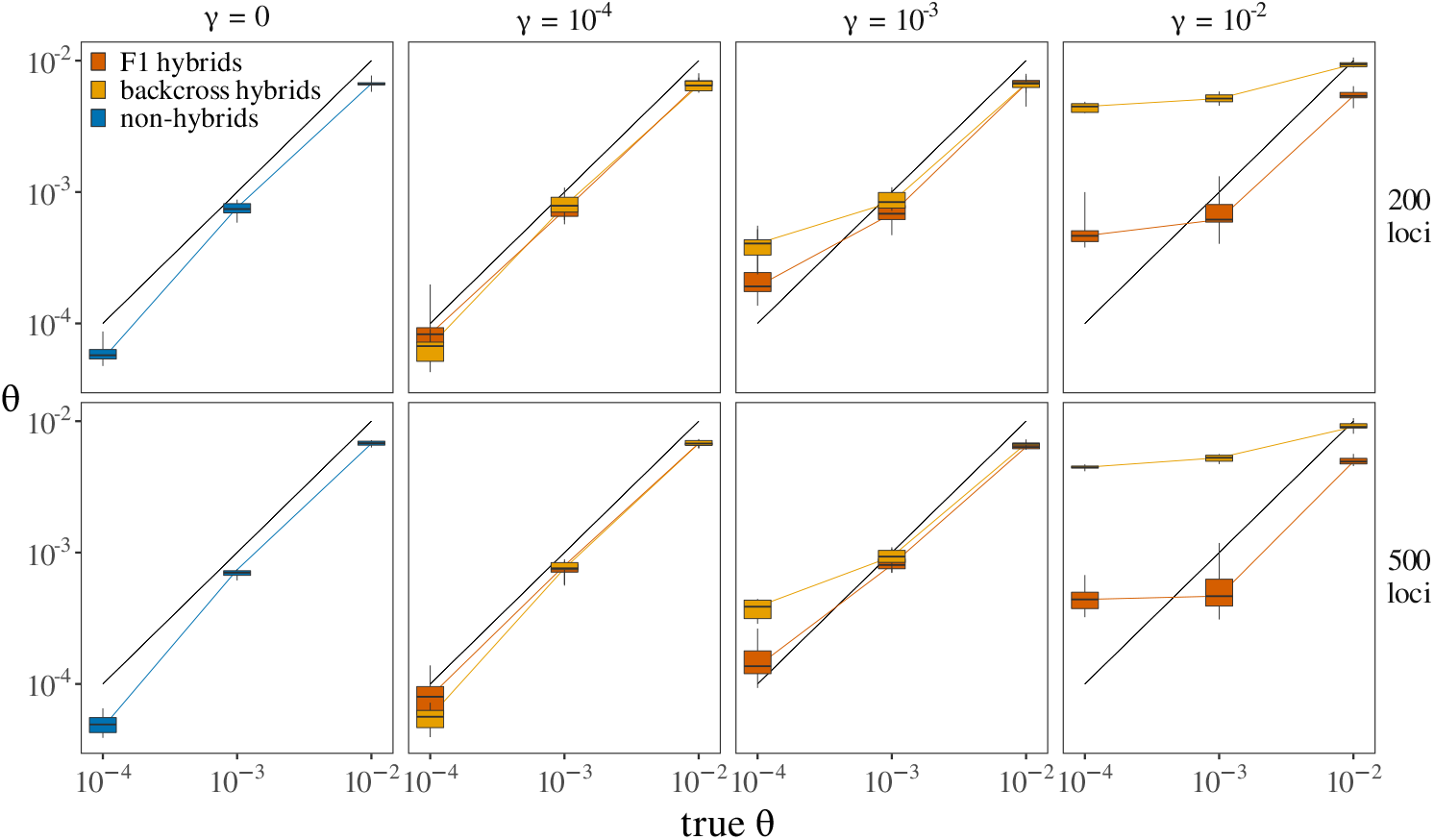
Ancestral population mutation rate estimation with varying number of loci. Each box represents the distribution of estimated *γ* values across 10 simulated individuals. Every individual consists of a collection of loci simulated under a given combination of true *θ* (given in x axis) and *γ* (given in headers) values. In non-hybrid individuals *γ* is always zero. In backcross hybrids, the true value of *γ* is that given in the header, but only for a binomial proportion of loci (as described in the main text). Rows give results obtained when varying loci set size.

**Figure S2:**
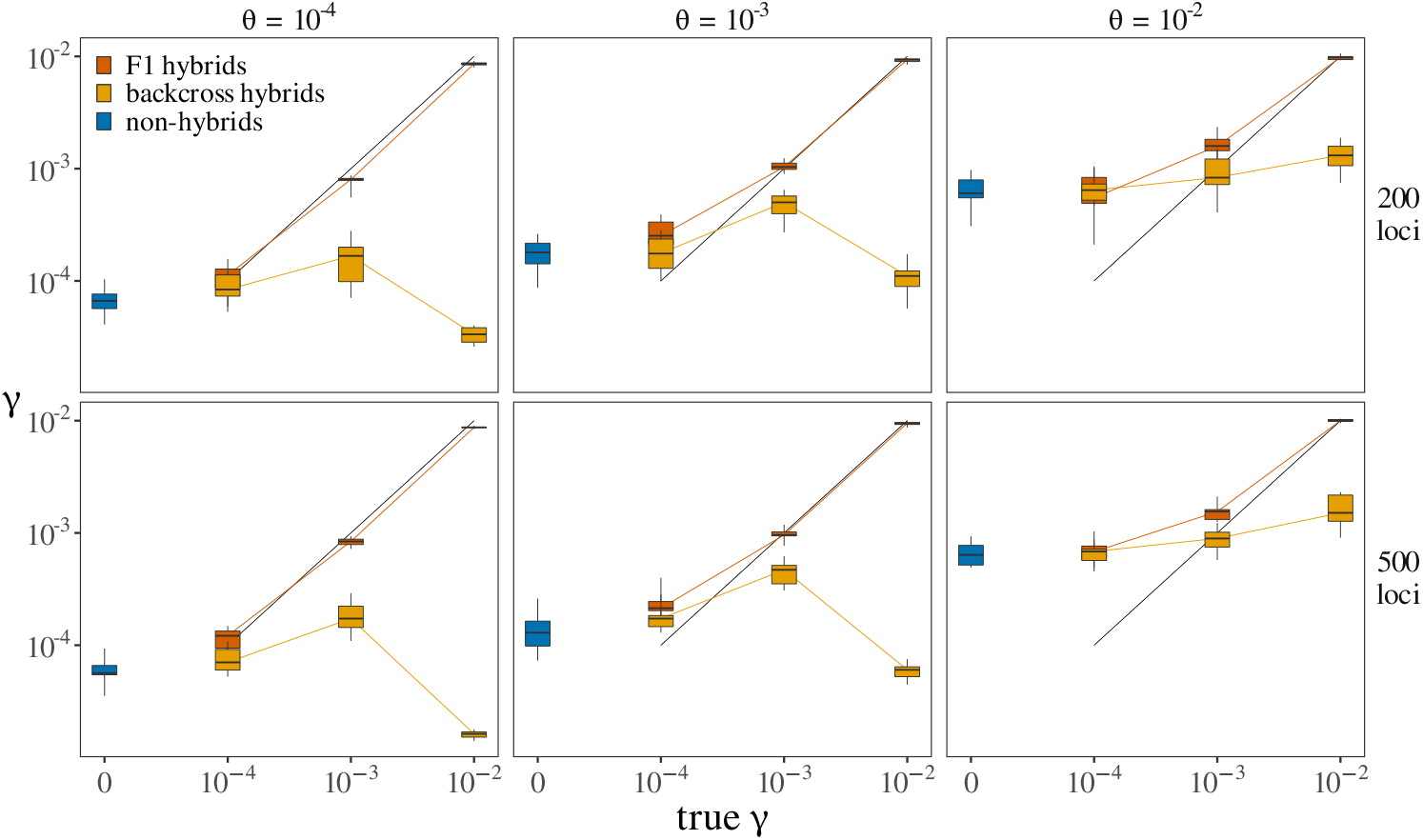
Divergence estimation with varying number of loci. Each box represents the distribution of estimated *γ* values across 10 simulated individuals. Every individual consists of a collection of loci simulated under a given combination of true *θ* (given in headers) and *γ* (given in x axis) values. In non-hybrid individuals *γ* is always zero. In backcross hybrids, the true value of *γ* is that given in the header, but only for a binomial proportion of loci (as described in the main text). Rows give results obtained when varying loci set size. The middle row is the same as the first row from figure 2 in main text.

**Table S4:**
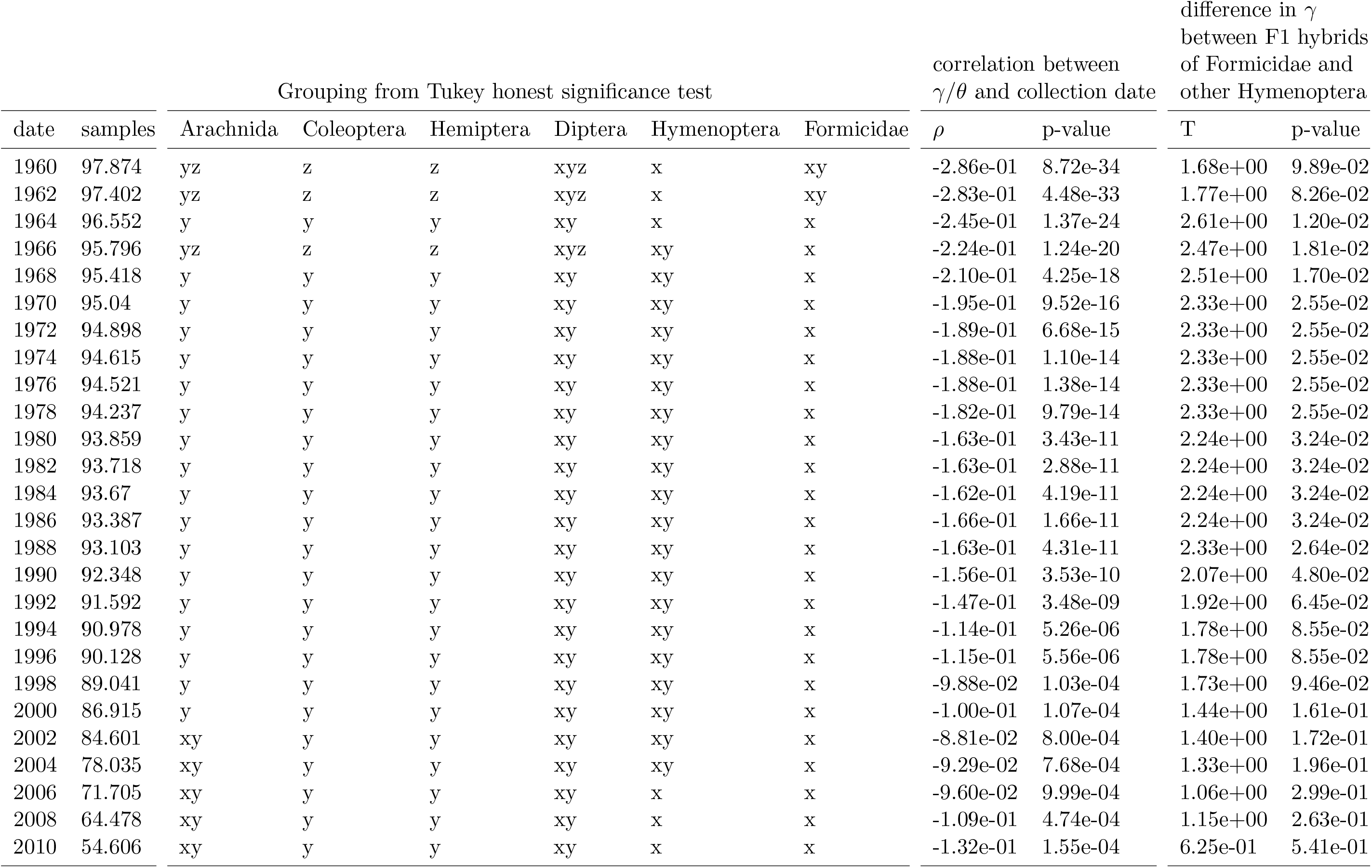
Main statistical results with varying maximum collection date. The table gives statistical results obtained when discarding all non-dated samples and all dated samples collected before a given threshold date (first column). Letters in columns 2 to *γ* summarize the result of a post-hoc Tukey honest significance test, carried out on *γ*/*θ* values across phylogenetic groups. The next column pair gives the pearson correlation coefficient between *γ*/*θ* and collection date. The last column pair summarizes results obtained when assessing the difference in mean *γ* between F1 hybrids (i.e., samples with *γ*/*θ* > 1) of Formicidae and non-Formicidae Hymenoptera with Student’s t-tests.

## Notes

https://zenodo.org/record/5415947

### Competing Interest Statement

The authors have declared no competing interest.

